# Hyperglycemic O-GlcNAc transferase activity drives cancer stem cell induction in TNBC

**DOI:** 10.1101/2022.03.14.484003

**Authors:** Saheed Ayodeji, Bin Bao, Emily A. Teslow, Lisa A. Polin, Greg Dyson, Aliccia Bollig-Fischer, Charlie Fehl

## Abstract

Enhanced glucose metabolism is a feature of almost all cancers, but downstream functional effects of aberrant glucose flux are difficult to mechanistically determine. The objective of this study is to characterize a mechanism by which elevated glucose level drives a tumorigenic pathway in triple negative breast cancer (TNBC). We used chemical biology methods to track how a metabolite of glucose, N-acetylglucosamine (GlcNAc), is linked to the transcriptional regulatory protein tet-methylcytosine dioxygenase 1 (TET1) as an O-linked GlcNAc post translational modification (O-GlcNAc). In this work, we revealed that intracellular protein glycosylation by O-GlcNAc is driven by high glucose levels in TNBC models, including on TET1. A single enzyme, O-GlcNAc transferase (OGT), is responsible for catalyzing protein modification of O-GlcNAc. We showed that OGT activity is higher in TNBC cell lines compared to non-tumor breast cell lines and is associated with hyperglycemia. Furthermore, enhanced OGT activity activated a pathway for cancer stem-like cell (CSC) reprogramming in TNBC cells. In our model, O-GlcNAcylated TET1 upregulated expression of splicing factor TAR-DNA binding protein (TARDBP), which drives CSC induction as well as higher OGT levels. We show that this OGT-TET1-TARDBP axis “feeds-forward” in hyperglycemic conditions both in cell lines and diet-induced obese mice, which displayed higher blood glucose levels and tumor O-GlcNAc levels than lean littermates. This data converges on a novel pathway whereby hyperglycemia drives aberrant OGT activity, activating a pathway for CSC induction in TNBC. Our findings partially explain a key aspect of how obesity is associated with TNBC risk and negative outcomes.

**Statement of Implication:** This work presents a novel mechanism to explain how obesity is a risk factor for triple-negative breast cancer via elevated sugar-transfer activity by O-GlcNAc transferase in hyperglycemia, leading to the induction of a cancer stem-like cells and revealing a targetable pathway in obesity-associated tumors.

## Introduction

Glucose use is higher in most cancers, typically associated with aberrant metabolism whereby glucose is rewired through aerobic glycolysis to create additional biosynthetic intermediates,^1,2^ the so-called “Warburg effect.” Breast cancers display the “Warburg Effect” very early in their progression.^3^ Breast cancer is the most common cancer in women and the leading cause of death.^4^ Among breast tumor subtypes, triple negative breast cancer (TNBC) is particularly devastating because it lacks treatment options normally available to breast cancer.^5^ A TNBC diagnosis means that the tumor cancer cells lack expression of three hormone receptors: the estrogen receptor ER*α*, the progesterone receptor and the epidermal growth factor family/receptor tyrosine kinase family member HER2.^5^ Because TNBC lacks hormone drivers, these tumors can occur earlier than other breast cancer subtypes including in pre-menopausal women.^6,7^ Underlying mechanisms of TNBC tumorigenesis are still under study, but a link between obesity and TNBC is noted.^8,9^ Obesity is an aberrant physiological state with features of hyperinsulinemia, chronic inflammation, and chronic hyperglycemia, which together offer us clues to look for associated risk factors.^10,11^ Along this line of evidence, we recently published a pathway that connects obesity-induced inflammation to TNBC tumorigenesis.^12,13^ Here, we add to our pathway through a converging mechanism for hyperglycemia to further enhance obesity-associated TNBC tumorigenesis.

One of the intracellular mechanisms affected by heightened glucose levels is protein modification through sugar linkages.^14-16^ In particular, the dynamic and reversible O-linked modification of serine and threonine residues with N-acetylglucosamine (O-GlcNAc) is driven by glucose.^15,17^ Roughly 2-5% of cellular glucose is converted by the hexosamine biosynthetic pathway to the donor sugar UDP-GlcNAc (uridine diphosphate-N-acetylglucosamine).^18^ There is a growing collection of associations between O-GlcNAc protein modifications and enhanced cancer progression,^19-21^ but our mechanistic understanding of O-GlcNAc-driven tumors remains far from solved because over 7000 proteins are known to be O-GlcNAc modified.^22,23^ Just one human enzyme is responsible for all protein O-GlcNAc modifications, O-GlcNAc transferase (OGT).^24^ This extreme reliance on a single protein for a cell-wide O-GlcNAc protein modifications makes OGT required for cell proliferation.^25^ Downstream effects of OGT have been observed in breast cancers, including activation of the oncogenic transcription factor FoxM1^26^ and the stem cell factor KLF8.^27^ Between breast cancer subtypes, OGT inhibition is most effective in TNBC.^28^ These studies suggest that OGT-driven cell events are intriguing biological targets for TNBC, especially if downstream pathways can be identified that differentially halt aberrant proliferation but maintain normal cell health.^29^

Here we show that OGT activates a tumorigenic pathway in TNBC through elevated OGT activity. In particular, direct O-GlcNAc modification of the protein tet methylcytosine dioxygenase 1 (TET1) activates our previously reported pathway^12,13^ leading to cancer stem-like cells (CSCs), the minor fraction of tumor cells that drive tumor initiation as well as re-initiation in metastasis.^30,31^ Furthermore, we reveal that hyperglycemic cell conditions not only enhance OGT activity but also its expression in TNBC cell lines and diet-induced obese mouse models. Our data suggest that this O-GlcNAc-driven pathway may be a targetable link between metabolic disorders and TNBC risk via this CSC pathway. Our data also suggests a potential mechanism for upregulated OGT levels in various cancers including TNBC.^19^

## Materials and Methods

### Cell Culture and Reagents

MDA-MB-231, MDA-MB-468, and HCC70 cells were originally purchased from ATCC. MCF 10A was obtained from Karmanos Cancer Institute (former Michigan Cancer Foundation). SUM 149 cells were gifted to us by Dr. Stephen Ether, whose lab developed the cell line. All cells were screened – free from mycoplasma using MycoAlert™ Mycoplasma Detection kit (LT0-118, Lonza group Ltd.) and MycoAlert™ control set (LT07-518, Lonza group Ltd.). The cells were all cultured in humidified sterile incubator conditioned at 5% CO_2_ at 37°C. MDA-MB-231 and MDA-MB-468 cells were grown in high glucose (4.5 g/L) or low glucose (1.0 g/L) DMEM media (ThermoFisher) supplemented with 10% Fetal Bovine Serum (FBS) and 1% Penicillin/Streptavidin (Pen/Strep). HCC70 cells were cultured in high glucose (4.5 g/L) or low glucose (1.0 g/L) RPMI-1640 media (ThermoFisher) supplemented with 10% FBS and 1% Pen/Strep. MCF 10A cells were cultured in a 1:1 DMEM: F-12 medium supplemented with 5% horse serum, 20 ng/mL EGF, 0.5 mg/mL Hydrocortisone, 100 ng/mL Cholera toxin and 10 μ/mL insulin. SUM 149 cells were cultured in Ham’s F-12 media (ThermoFisher) supplemented with 10% FBS and 1% Pen/Strep. We validated all cell lines employing short tandem repeat analyses using PowerPlex® 16 system (Promega).

### Chemical Synthesis and Chemoenzymatic labeling

Chemical probes were synthesized according to published reports, including OGA inhibitor Thiamet-G^32^ and azide labeled O-GlcNAc metabolic labeling sugar Ac_4_GalNAz^33^ were synthesized, purified, and characterized using ^1^H-NMR, ^13^C-NMR, and mass spectrometry to match reported spectra. OGT inhibitor OSMI-4^34^ was originally provided by the Suzanne Walker group at Harvard Medical School, and then purchased from GLP Bioscience (Cat# GC31517). Inhibitors were solubilized in DMSO. Click-IT™ O-GlcNAc enzymatic labeling kit (Cat#C33368) was obtained from ThermoFisher. Biotin-alkyne (#1266-5) and TAMRA-alkyne (#1255-5) were purchased from Click Chemistry Tools (Scottsdale, AZ). O-GlcNAc Chemoenzymatic labeling using the Click-IT™ kit were carried out as recommended by the manufacturer (ThermoFisher). Each labeling experiment was started with 200 μg protein lysate (per sample) from cell line cultured in Low (1.0 g/L) or High (4.5 g/L) Glucose media for 72 hours. The labelled lysate was then either “clicked” on with TAMRA-alkyne or biotin-alkyne using copper catalyzed azide-alkyne cycloaddition (CuAAC) click chemistry reaction protocol supplied by ThermoFisher. Biotin-labelled samples were then immunoblotted and visualized with Streptavidin-HRP antibody on an iBright™ FL-1500 imaging system (ThermoFisher).

### Western blotting

Western blotting procedures were carried out following standard protocol. Briefly, cells were allowed to reach 80 – 90% confluency and collected using RIPA buffer (ThermoFisher, #89900) or 1% NP-40 buffer (150 mM NaCl, 50 mM Tris-Cl (pH 7.4), 1% NP-40) supplemented with Roche cOmplete™ protease inhibitor cocktail (#11836170001). Protein lysates from OSMI-4-treated cells were collected after 72 hours of probe treatment. Cell lysates were clarified by centrifugation at 17,000 x g for 15 minutes at 4°C. The supernatant containing soluble protein fractions were collected and quantified using Pierce™ Rapid Gold BCA Protein Assay kit (#A53225) and analyzed by SDS-PAGE and protein specific antibody western blot. Antibodies used for Western blot are: Anti-OGT (CST, #24083), Anti-O-GlcNAc MultiMAb (CST, #82332), Anti-Actin (Sigma, #A3853), Anti-Tubulin (Invitrogen, #MA1-80017), Anti-TET1 (Abnova Taiwan, #H00080312), Anti-TARDBP (Abnova, #H000234350-M01)

### Transient Knockdown

For OGT knockdown, we transfected ON-TARGET plus SMART pool human OGT siRNA ((Dharmacon™ # L-019111-00-0005) and ON-TARGET control pool non-targeting pool siRNA (#D001810-10-05) as control using DharmaFECT™ transfection reagent as described by the manufacturer. The SMART pool is a combination of 4 different siRNA oligos optimized for knockdown in human cell lines, which we used here in place of two distinct siRNA sequences for OGT knockdown and its corresponding SMART pool control knockdown. For TARDBP knockdown, we transfected two siRNA (ThermoFisher, #s23829 and #s530935) targeting TARDBP mRNA and *silencer*^*TM*^ select negative control siRNA (ThermoFisher, #4390846).

### Semiquantitative real-time RT-PCR (qRT-PCR)

RNA was isolated from cultured and treated cell lines or from pulverized snap-frozen tumor tissue using Qiagen RNeasy kit following the manufacturer’s procedure. mRNA samples were collected from OSMI-4-treated cells after 24 hours. The RNA was converted to cDNA using the High-Capacity RNA-to-cDNA kit (ThermoFisher). The cDNA was diluted to 50 ng/μL working concentration for all samples and 2 μL of the working cDNA samples were then combined with SYBR Green PCR Master Mix reagents (ThermoFisher) to make up 20μL reaction volumes in 96-wells plates. qRT-PCR reactions were run in three technical replicates using StepOnePlus Real-Time PCR System (ThermoFisher). mRNA expression of PUM1 was used as control for all qRT-PCR experiments and the expression data were calculated using the delta-delta Ct method.

### Animal work

All experiments and procedures involving animals and their care were pre-reviewed and approved by the Wayne State University Institutional Animal Care and Use Committee. All tumor samples analyzed in this work had been previously prepared and snap-frozen during the data collection for the prior study.^12^

### Statistical analysis and bioinformatics

Statistical analysis was performed using Bioconductor R 3.3.2 and GraphPad Prism. Graphs were generated with GraphPad Prism. *P* values ≤ 0.05 are reported as significant and \**P* ≤ 0.05, \*\**P* < 0.01, \*\*\**P* < 0.001 indicates level of significance. Linear regression or Student’s t-test (two-sided) were applied when two conditions were compared, including qRT-PCR analysis. For analysis of 3 or more experiment conditions, a mixed-model approach was applied. To identify significant differences among gene expression data from The Cancer Genome Atlas (TCGA), analysis of variance (ANOVA) with Tukey’s HSD test was performed. For analysis of gene expression according to intrinsic subtyping, microarray data and PAM50 subtype annotation from TCGA were downloaded via cBioPortal. For analysis of clinically relevant subtypes, RNA-sequencing data from TCGA were downloaded from The National Cancer Institute Genomic Data Commons data portal.

## Results

### OGT is overexpressed in tumors from TNBC patients and cell lines vs. tumors from other types of breast cancer

O-GlcNAc transferase (OGT) is the single gene product responsible for all O-linked modification of protein serine/threonine residues with the monosaccharide N-acetylglucosamine (GlcNAc).^24^ OGT is an essential gene in all cell types, underscored by it being the least frequently mutated glycosyltransferase (GT) in humans.^25^ Despite the fact that OGT is expressed and its levels are tightly regulated in all human cells,^35^ our analysis of patient data in The Cancer Genome Atlas (TCGA) found an significant elevation of OGT in TNBC tumors over normal matched breast tissue. Compared to normal tissue, OGT is significantly overexpressed in TNBC patient tumors (**Fig 1A**). Between subtypes of breast cancer tumors, including HER2-enriched and estrogen/progesterone receptor-expressing luminal breast tumors, TNBC showed significantly higher OGT expression. (**Fig. 1B**).

**Figure 1:**
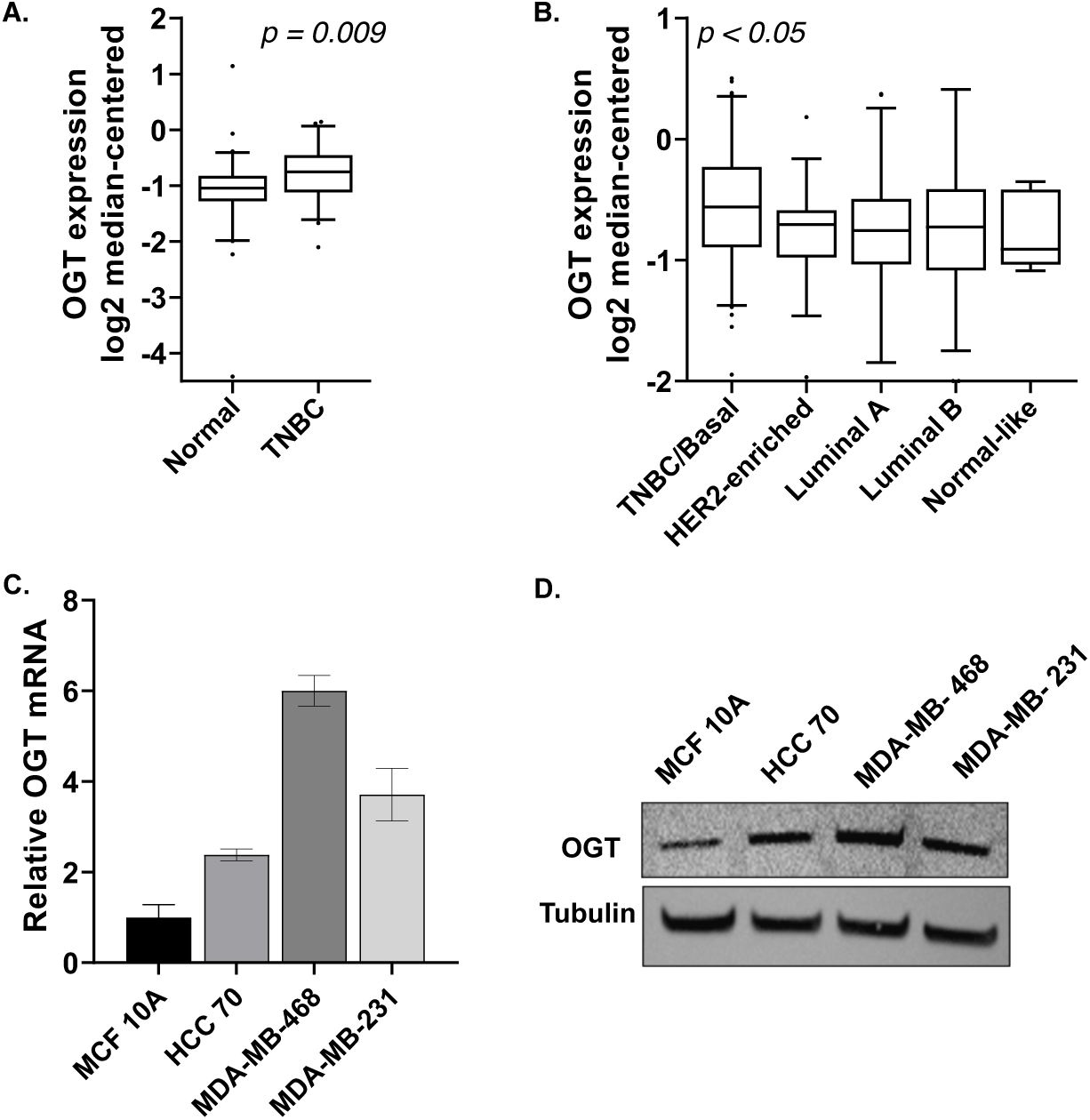
OGT is upregulated in TNBC tumors and cell lines. **A-B**) Whole genome expression analyses of human patient samples targeting OGT mRNA expression in TNBC, matched breast tissue, and other subtypes of breast cancer. Line equals median, means were compared. **C**) qRT-PCR and western blot analysis of OGT in each of four TNBC cell lines. Bars, SEM 3 independent experiments. **D**) Western blot analysis of OGT in non-tumor breast cell line MCF 10A and three TNBC cell lines.

We analyzed OGT levels in three established TNBC cell lines (HCC70, MDA-MB-468, MDA-MB-231) as compared to the non-cancer breast epithelial cell line MCF10A (**Fig 1C-D)**. We observe different levels of basal OGT expression between the three TNBC lines in our panel, but in all cases, OGT levels were upregulated. Analyzing OGT characteristics across these three cell lines approaches better representation of heterogeneous patient populations.

### OGT catalytic activity drives an obesity-driven cancer stem-like cell induction pathway

As outlined in the introduction, we recently characterized a pathway the connects obesity phenotypes with elevated TNBC tumorigenesis via CSC induction.^12,13^ More importantly, inhibiting this pathway targets TNBC CSC cells directly.^36^ Briefly, in TNBC cells, activity of the enzyme tet methylcytosine dioxygenase 1 (TET1) enables overexpression of TAR DNA-binding protein (TARDBP), a nucleotide binding protein. Elevated TARDBP activity positively regulates another splicing factor, serine and arginine rich splicing factor 2 (SRSF2). Higher TARDBP and SFRSF2 levels produce the tumorigenic “splice variant 2” of methyl CpG-binding protein 2 (MBD2_v2). MBD2_v2 leads to the stem cell reprograming factor NANOG and CSC expansion (**Fig 2B**). This pathway was upregulated in diet induced obesity (DIO) mouse models vs. lean littermates, which we attributed to elevated TET1 activity due to H_2_O_2_ levels in obesity-driven inflammation.^13^

**Figure 2:**
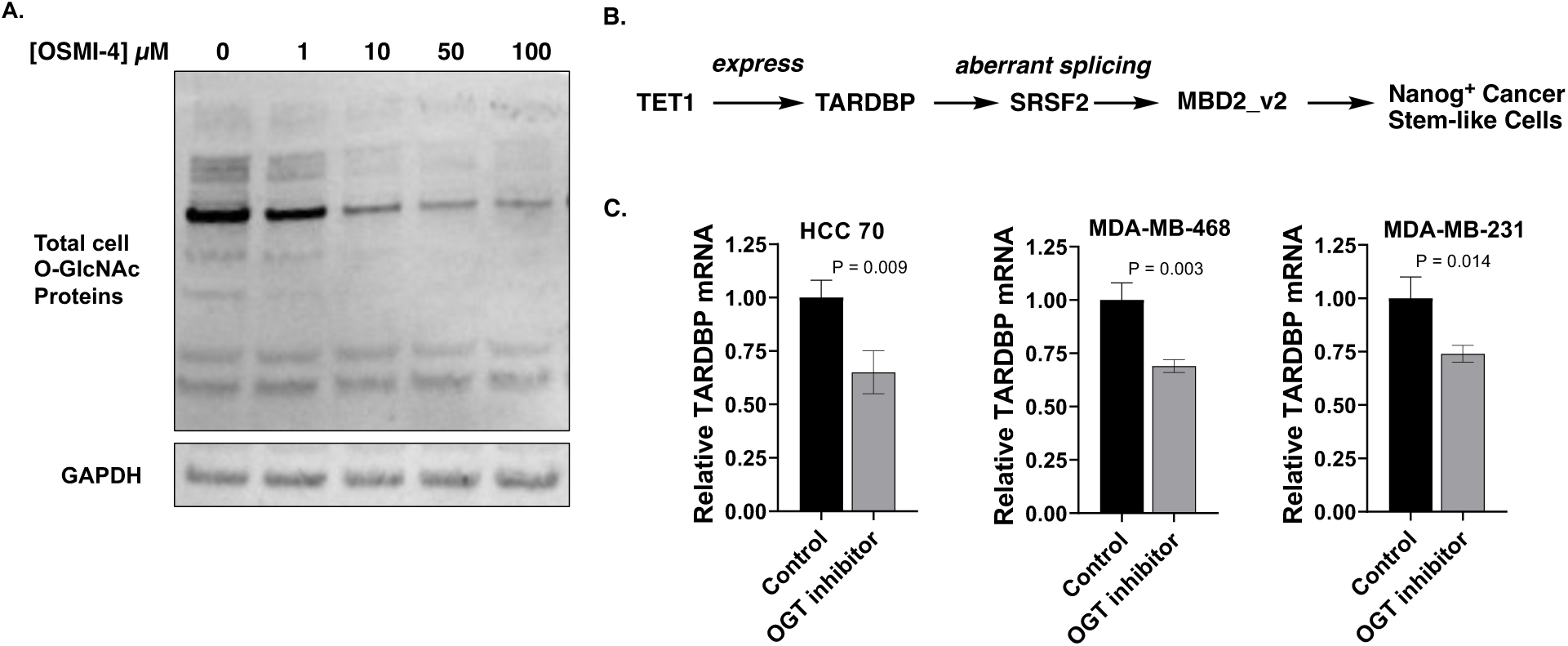
OGT inhibition reduces global O-GlcNAcylation, including TET1 activity. **A**) Dose-response curve for total cellular O-GlcNAcylation with OSMI-4 in MDA-MB-468 cells. Western blot performed with anti-O-GlcNAc MultiMAb. **B**) Our published mechanism for a TET1-driven pathway to cancer stem-like cells in TNBC. **C**) TARDBP mRNA analysis by qRT-PCR upon OGT inhibition with OSMI-4-(10 μM, 24 hours). Data is normalized to vehicle-treated controls. Bars, SEM 3 experiments. N = 3.

Intriguingly, we noted that TET1 is reported to have tight affinity for a protein-protein interaction with OGT.^37,38^ O-GlcNAcylation of TET1 by OGT activates its catalytic activity for 5-hydroxymethylcytosine formation in embryonic stem cells, leading to enhanced target gene expression.^39,40^ We hypothesized that elevated OGT levels and activity in TNBC cells may similarly drive this pathway in cancer stem cells. We therefore sought to test the ability of OGT to regulate TET1 and the following CSC induction pathway. To measure TET1 activity, the downstream target gene TARDBP was chosen for qRT-PCR analysis. A small molecule probe, OSMI-4, was recently reported to inhibit OGT with a *K*_i_ of 8 nM and high selectivity over other human glycosyltransferases.^34,41^ We treated TNBC cell lines with the OGT inhibitor OSMI-4 and observed globally inhibited overall O-GlcNAcylation, as measured with a pan-O-GlcNAc antibody to detect global cell O-GlcNAc modifications on proteins (**Fig 2A**). Active TET1 leads to TARDBP expression, so we used TARDBP mRNA levels as a readout for TET1 activity. Treatment with OSMI-4 revealed that inhibition of OGT led to reduced TARDBP expression levels in TNBC cell lines (**Fig 2C**).

Next, we used a chemical strategy to label O-GlcNAc modifications on proteins in TNBC cells (**Fig 3A**). Briefly, an engineered enzyme that installs an azide chemical label directly onto the O-GlcNAc modification on proteins was used to “tag” O-GlcNAcylated proteins in TNBC cell lysates.^42,43^ The resulting azide-labeled proteins can be directly visualized with copper-catalyzed bioorthogonal “click” chemistry, which drives an irreversible, copper-catalyzed azide-alkyne cycloaddition (CuAAC) reaction between the O-GlcNAc-azide species and a fluorescent alkyne. These chemical biology labeling reactions revealed the total O-GlcNAcylation patterns in TNBC cells, which were compared using total fluorescence counts (**Fig 3B**). Click reactions with biotin-alkyne installed an affinity enrichment handle that enabled selective protein visualization by western blot following streptavidin enrichment, i.e. using antibodies against TET1. We used this “biotin click/enrichment/TET1 immunoblot” sequence to verify that TET1 is indeed O-GlcNAc modified in TNBC cells. Moreover, cells grown in high vs. low glucose show elevated TET1-O-GlcNAcylation, indicating that the O-GlcNAc modification increase is glucose-driven, with TET1-O-GlcNAc levels increasing by at least 2-fold (**Fig 3C**). As a final confirmation of TET1 O-GlcNAcylation, we overexpressed tagged-TET1 catalytic domain and used metabolic labeling to install an O-GlcNAc chemical reporter on TET1 in HEK293T cells (**Supplemental Figure 1**).^33^

**Figure 3:**
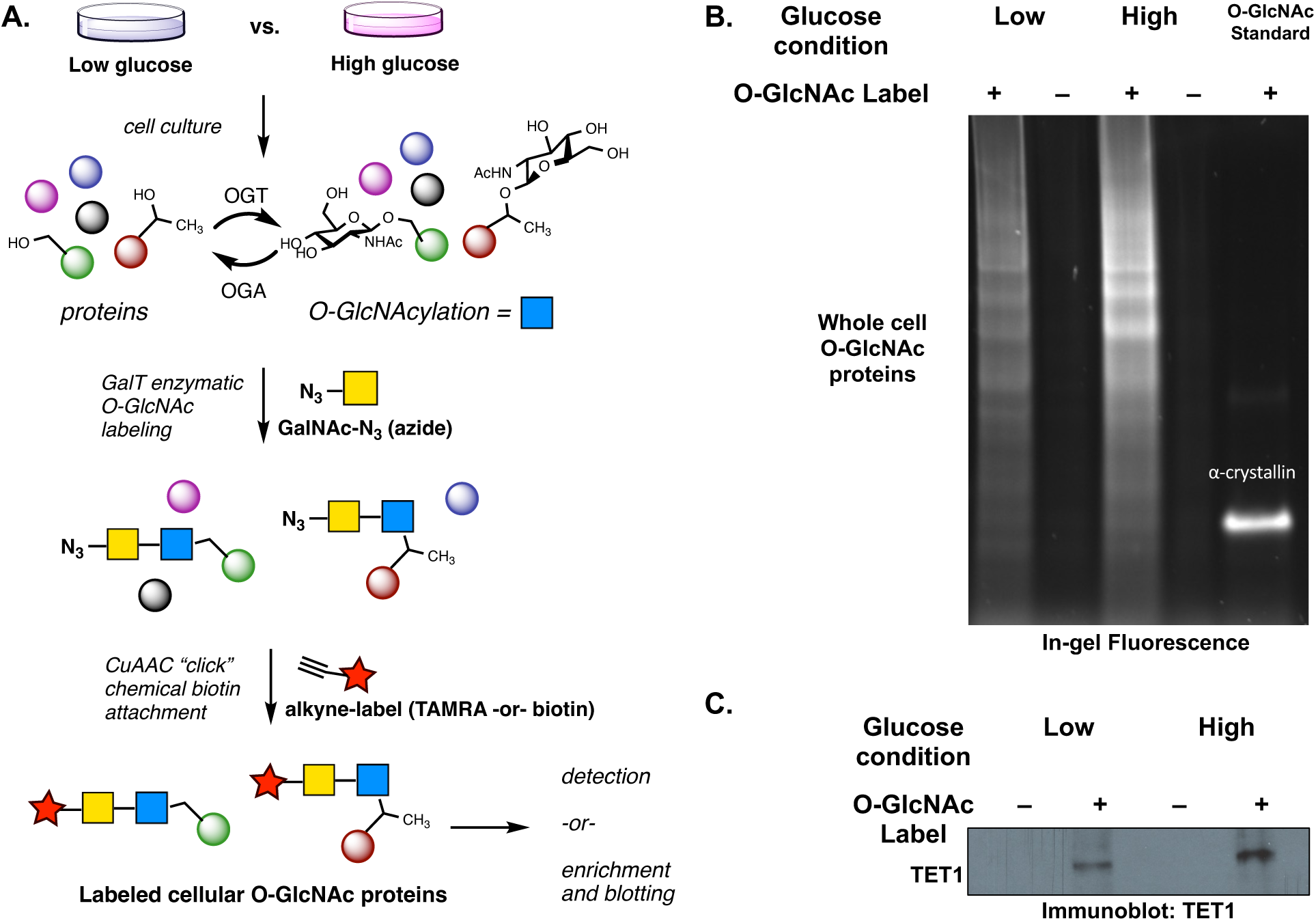
Chemical labeling of O-GlcNAc reveals TET1-O-GlcNAc modifications in TNBC cells. **A**) Total cellular O-GlcNAc detection scheme: chemoenzymatic labeling using GalT (N-acetylgalactosaminyl transferase mutant Y189L) and copper-catalyzed azide-alkyne cycloaddition (CuAAC) “click” chemistry labeling of O-GlcNAc sites. **B**) Total cell O-GlcNAcylation measured with TAMRA-red fluorophore-alkyne click reaction. Alpha-crystallin was used as a known O-GlcNAcylated positive control protein. The labeling was normalized by using 200 ug of total protein input for all reactions. **C**) Total O-GlcNAc enrichment using biotin “click” labeling and streptavidin beads followed by TET1 Western blot. High glucose (4.5 g/L) and low glucose (1.0 g/L) Dulbecco’s Modified Eagle Medium (DMEM) was used in these experiments. N = 2, representative images shown.

Taken together, the OGT inhibition study and direct chemical biology labeling study show that TET1 is an OGT substrate in TNBC cells. Moreover, reducing TET1 O-GlcNAc modification via OGT inhibition revealed less TARDBP production. We hypothesized that suppressing OGT would inhibit our reported CSC-inducing tumorigenic pathway because OGT activity regulates TET1.^13^

### Pathway engineering confirms that OGT is involved in and regulated by a CSC pathway

We have established that TET1,^13^ SRSF2,^12^ TARDBP,^13^ and MBD2_v2^44^ elements are all necessary for TET1-driven CSC reprogramming as part of a linked pathway (summarized in **Fig 2B**) using knockdown and overexpression. Here, we used knockdown studies to engineer TNBC cell lines to verify OGT effects and possibly regulation as part of this TET1-driven pathway. OGT knockdown recapitulated our inhibition results with OSMI-4 inhibitor (**Fig 4A-B**). The combined data from OGT knockdown and inhibitor treatment confirm that OGT catalytic activity regulates TET1 because OSMI-4 is known to inhibit OGT catalytic activity through competitive substrate binding.^34^

**Figure 4:**
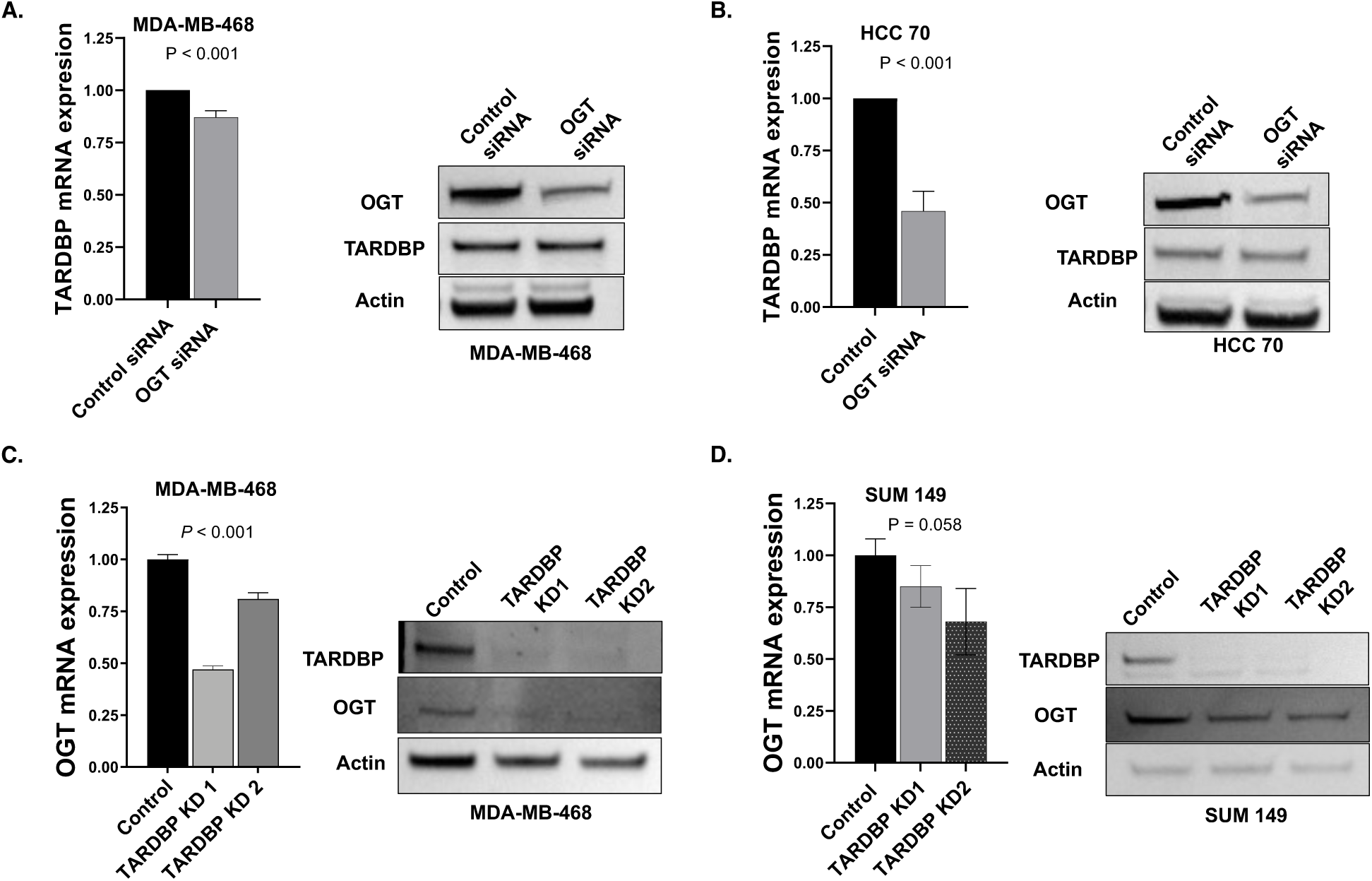
Knockdown studies confirm the pathway organization. **A-B**) TNBC cells were treated with SmartPool OGT siRNA (4 targeting sequences) or a control, non-targeting SmartPool siRNA for 48 h. TARDBP mRNA and protein levels were measured. **C-D**) TNBC cells were treated with two different TARDBP siRNA sequences and OGT mRNA and protein levels were measured. Data is normalized to non-silencing siRNA-treated controls. Bars, SEM 3 experiments. N = 3 for all experiments.

We also knocked down or inhibited downstream pathway members and analyzed OGT expression levels. Surprisingly, we discovered that TARDBP is necessary for high OGT expression levels despite being downstream in our proposed pathway (**Fig 4C-D**, also see **Fig 2B**). A known feature of OGT expression level regulation is through alternative splicing to produce mRNA encoding the functional gene product or, via use of a “detained intron,” an alternative transcript that is rapidly degraded via nonsense-mediated mRNA decay.^35^ Cells use detained intron splicing mechanisms to rapidly regulate OGT levels to maintain homeostasis. Because TARDBP is a DNA binding protein and a splicing modulator,^45^ we were able to use the SpliceAid database^46^ to reveal a canonical TARDBP binding site on OGT mRNA (**Supplemental Figure 2**). A potential mechanism for TARDBP regulation of OGT, despite being downstream in the CSC pathway we propose, is through TARDBP’s predicted OGT mRNA binding activity.^45^ We note differences in the level of OGT reduction upon TARDBP knockdown between different cell lines, potentially reflecting differences in total cell OGT mRNA levels (see **Fig 1C-D**).

Upon OGT knockdown or inhibition (see **Fig 2C**, above), TARDBP protein levels are reduced but not completely abolished. Our OGT suppression data is consistent with our hypothesis that OGT binds and activates TET1 activity via O-GlcNAcylation, as observed in embryonic stem cells.^38,39^ The effect of TARDBP knockdown and OGT reduction suggests that the pathway we propose additionally includes a reciprocal element of OGT regulation by TARDBP, likely via mRNA binding (**Supplemental Figure 2**). These data suggest that OGT may drive TARDBP levels, which allows further stabilization of OGT in a “feed-forward” manner (see **Discussion**).

### OGT is regulated by hyperglycemic cell conditions and in models of metabolic disease

If our hypothesis that OGT activity drives the level of TARDBP and thus stabilizes more OGT is true, we predicted that hyperglycemia would activate the pathway via elevated OGT substrate levels. To test this prediction, we characterized OGT and OGT-driven TET1 activity in high vs. low glucose media using chemical O-GlcNAc labeling to measure OGT activity. First, we used chemoenzymatic labeling to directly compare O-GlcNAc modifications on TNBC cells grown in high vs. low glucose media, as shown in **Fig 3** above for the HCC 70 model of TNBC. Indeed, two additional TNBC cell lines, MDA-MB-231 and MDA-MB-468, showed elevated total O-GlcNAc levels in high glucose media as well as higher specific O-GlcNAcylation of TET1 (**Fig 5A-B**).

**Figure 5:**
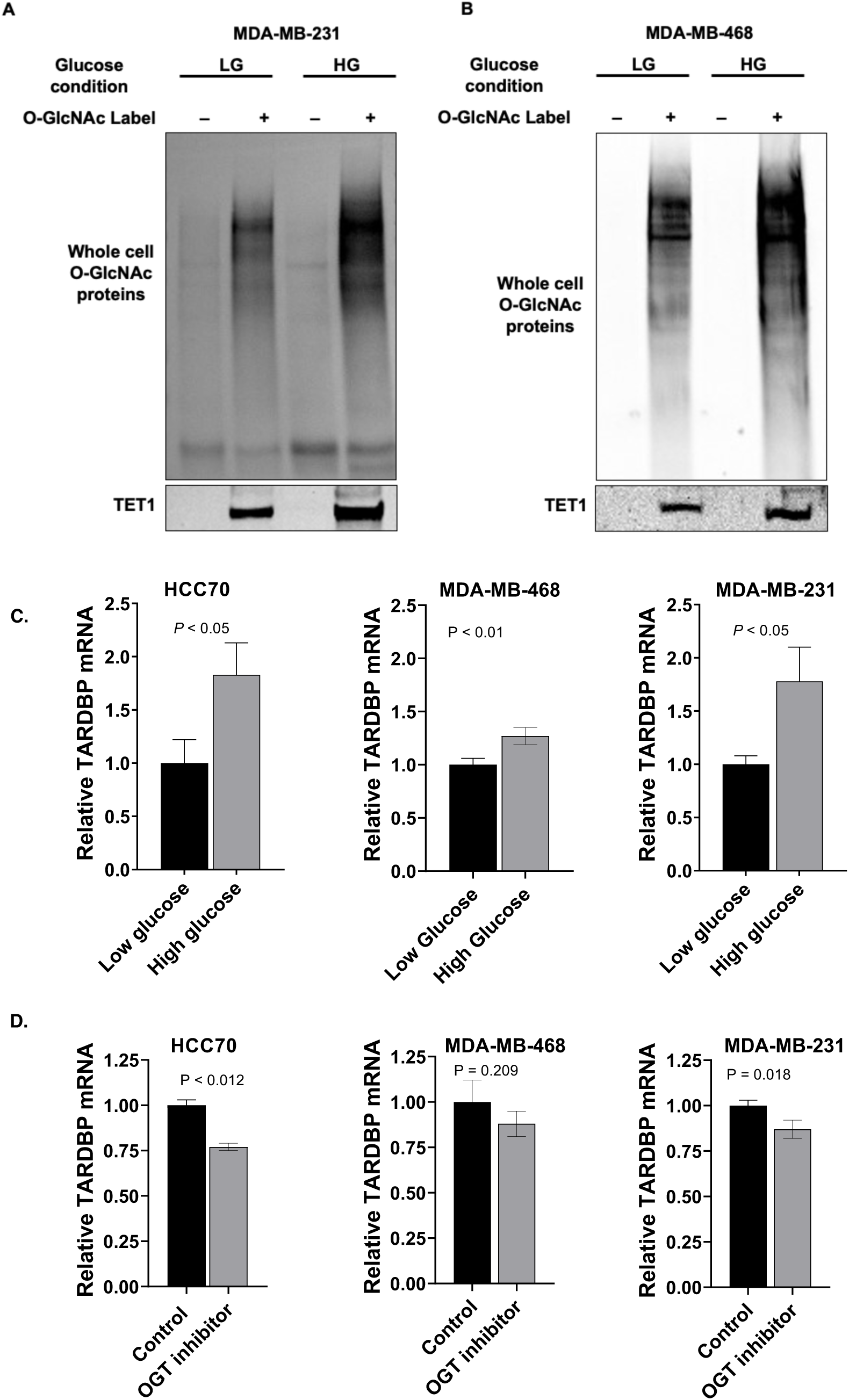
Chemoenzymatic labeling of additional TNBC lines reveal glucose-driven O-GlcNAcylation. **A-B**) Biotin “click” on O-GlcNAc modifications followed by total O-GlcNAc detection (blot: streptavidin, top panel). Streptavidin enrichment followed by TET1 blot showed O-GlcNAc levels on TET1 protein (IP: O-GlcNAc-biotin, blot: anti-TET1) under high glucose vs. low glucose conditions. **C**) Comparison of TARDBP mRNA levels by qRT-PCR between high and low glucose conditions. **D**) TARDBP mRNA levels in OSMI-4 treated (10 μM, 24 hours) TNBC cells in high glucose conditions were measured by qRT-PCR, normalized to non-treated controls. Bars, SEM 3 experiments, N = 3. High glucose = 4.5 g/L Low Glucose = 1.0 g/L

Consistent with evidence that OGT-catalyzed O-GlcNAcylation impacted TET1 activity, TET1 target TARDBP expression also increased when TNBC cells were exposed to high glucose media (**Fig 5C**). Furthermore, though TNBC cell lines still respond to OGT inhibition using OSMI-4, the inhibitory effect of reduced OGT activity on TARDBP production is reduced in significance (compare **Fig 2C** in low glucose with **Fig 5D**, high glucose).

In our previous reports, we find that diet-induced obesity (DIO) can upregulate pathway members TET1 and MBD2_v2 levels.^12,13^ We used B6.Rag1 knockout mice, an obesity-compatible mouse model,^47^ which gained weight and attained hyperglycemia after 5 weeks on high fat diet (**Supplemental Figure 3A-B**). We next analyzed OGT mRNA from TNBC tumors implanted in lean vs. DIO mouse models using qRT-PCR (**Fig 6A**). We saw significant OGT overexpression in TNBC tumors grown in DIO mice vs. lean littermates, suggesting that obesity can elevate OGT expression in tumors. We hypothesize that this effect operates through the OGT/TET1/TARDBP loop that, via enhanced TARDBP activity, “feeds-forward” to lead to higher OGT levels.

**Figure 6:**
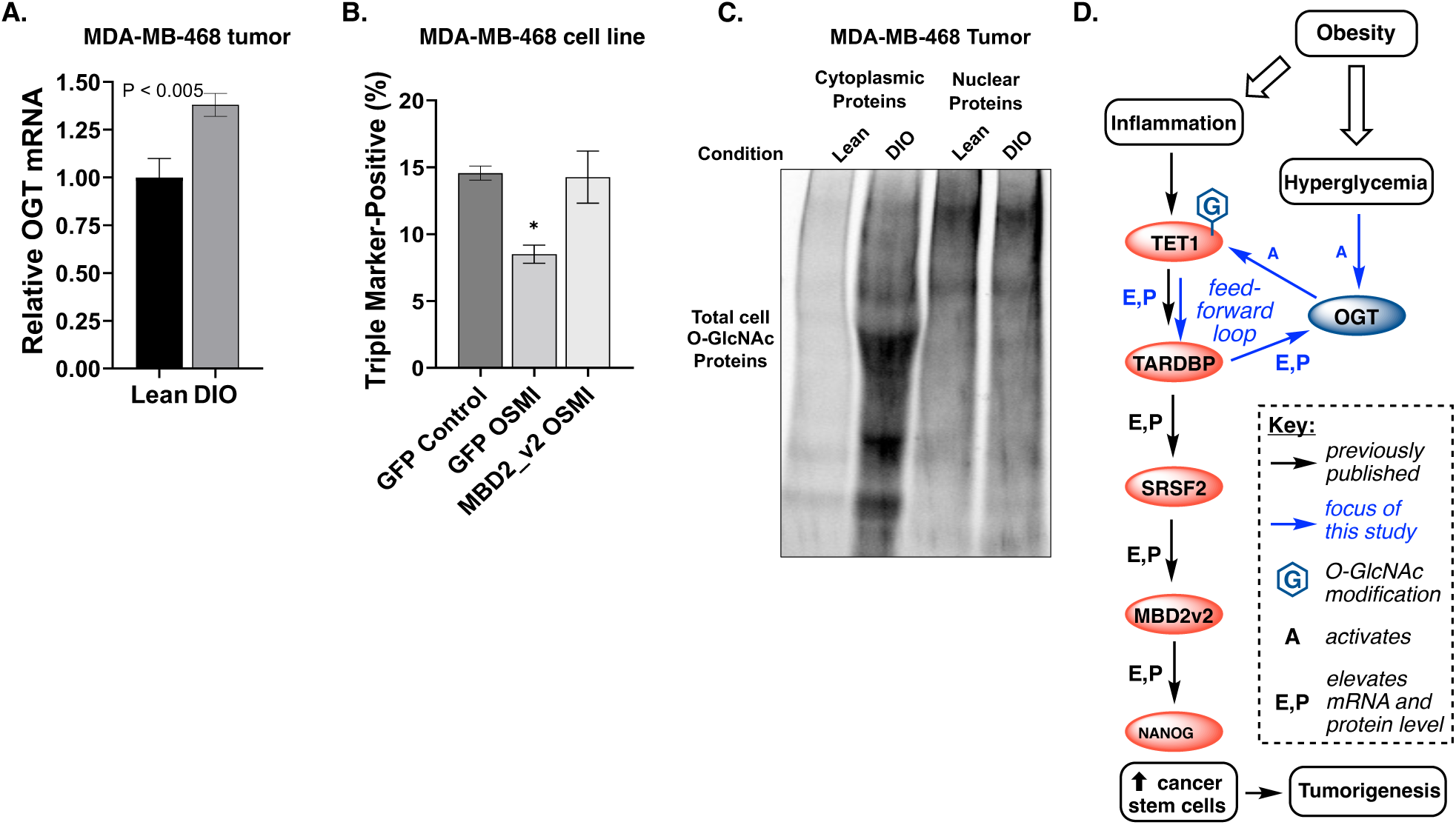
OGT and obesity-associated hyperglycemia-driven pathway to TNBC cancer stem induction. **A**) Mice fed a high-fat diet were allowed to attain diet-induced obesity (DIO) before tumor extraction and qRT-PCR. OGT levels in TNBC cell line MDA-MB-468-derived tumors harvested from DIO (N = 3) and lean control mice (N = 3). qRT-PCR analysis of each sample was performed in triplicate and the means normalized. **B**) Flow cytometry for triple stem cell marker expression under OGT inhibition. GFP indicates control cells and MBD2_v2 indicates MBD2_v2 overexpression (rescue). **C**) Chemoenzymatic labeling of total O-GlcNAc levels in TNBC tumors isolated from DIO vs. lean mice. N = 1. Nuclear and cytoplasm were extracted as detailed in Methods. **D**) Working Model for the OGT-driven pathway presented in this study.

To test whether TNBC cancer stem cell populations respond to OGT inhibition, we analyzed the production of stem cell markers in MBA-MB-468 cell lines.^36^ To test the effects of OSMI-4 on CSCs vs. non-CSC bulk cancer cells, we used fluorescence activated cell sorting (FACS) to analyze the population of CSCs using CD44, EpCAM, and CD133 sorting as “triple markers” of cancer stem-like cells.^48^ Treatment of GFP-vector-overexpressing TNBC cells with OGT inhibitors significantly depleted the population of CSCs by almost 50% (**Fig 6B**). The specific regulation of this pathway by OGT was confirmed by overexpression of MBD2_v2, an obesity related gene that is downstream of OGT in the CSC pathway presented here (see **Fig 2B**). MBD2_v2 overexpression was able to rescue OGT inhibition back to at least the same level as non-treated TNBC cells, confirming the OGT’s catalytic activity regulates this TET1-dependent CSC promoting pathway. Full flow cytometry data is shown in **Supplemental Figure 4**.

We also measured the impact of obese conditions on O-GlcNAc levels in tumors grown in DIO vs. lean mice using chemoenzymatic labeling (**Fig 6C**). Both cytosolic and nuclear fractions of a DIO-grown TNBC tumor compared with a lean TNBC tumor showed elevated overall O-GlcNAcylation. We did not isolate enough tumor material to enable TET1 pulldown studies, but our results mimic our *in vitro* high vs. low glucose cell culture conditions that elevate O-GlcNAc levels, including on TET1 protein.

Taken together, our data suggest that obesity and potentially other hyperglycemic diseases can both elevate OGT levels and drive a tumorigenic pathway in TNBC via the Working Model shown in **Figure 6D**. In this model, hyperglycemia activates OGT activity, creating a potentially dangerous situation in terms of the tumorigenic pathway under study because a majority of TNBC patients are overweight or obese at the time of diagnosis.^49,50^

## Discussion

Correlative studies between TNBC incidence and metabolic diseases including type 2 diabetes and obesity reveal that ca. 70% of TNBC patients are obese or overweight at the time of diagnosis.^51^ One aspect of this link could be metabolic reprograming in TNBC tumors.^52^ OGT serves as a key nutrient and metabolism sensor in human cells because its substrate UDP-GlcNAc is glucose-derived via the hexosamine biosynthetic pathway.^53^ Notably, OGT’s nutrient sensing role has been previously linked with breast cancer tumorigenesis via the transcription factor FoxM1, but since FoxM1 is not an OGT substrate there remains a missing link in this pathway.^26^ Establishing a mechanism for OGT-driven tumorigenesis would provide therapeutic insight into metabolic disease and TNBC risk. The reported physical interaction of OGT and TET1^38,39^ led us to hypothesize that our recently reported TET1 pathway to TNBC tumorigenesis^12,13^ might be regulated by OGT.

Altogether, our data led us to propose a novel working model for OGT overexpression in TNBC tumors and its impact on a tumorigenic pathway (**Fig 6D**). We previously demonstrated that obese physiology fuels a TET1-driven pathway leading to enhanced TNBC tumorgenicity.^12,13^ Here, we conclusively demonstrate that OGT is an upstream regulator of this pathway in a catalytic fashion; is itself regulated by splicing factors in this tumorigenic pathway; and that a second hallmark of obesity—hyperglycemia—further activates this pathway. To our knowledge, this study is the first demonstration that two distinct physiological hallmarks of obesity, systemic hyperglycemia and chronic inflammation, converge on an OGT and TET1-driven pathway to drive cancer stem-like cell induction. Rescue of OGT inhibition was performed by MBD2_v2 overexpression, further solidifying the upstream regulatory placement of OGT in this pathway. Reinforcing these *in vitro* results, our models of TNBC tumors in DIO mice show elevated OGT levels, O-GlcNAc levels, and support our published pathway whereby DIO mice display higher TNBC tumor initiation events than their lean littermates.^12^ These data provide a possible mechanism for the correlation between overweight and obese patients and TNBC diagnosis.

The unexpected regulation of OGT in this OGT->TET1->TARDBP pathway offers a potential mechanism to explain the higher levels of OGT mRNA that we observed in tumors between lean and DIO mice. In our model, upregulated tumor OGT levels could be a product of a pathway that “feeds-forward” in three steps. In first step, obesity triggers systemic hyperglycemia, which activates TET1 to produce higher levels of the DNA binding protein TARDBP. Second, TARDBP stabilizes or produces additional active OGT via splicing regulation—a known regulatory manifold for OGT^35^—to activate more TET1 -> TARDBP. Third, this process continues, resulting in the elevated OGT levels observed in TNBC patient data (**Fig 1A-B**) and our DIO mouse data (**Fig 6C**). This combined cellular and animal data suggest that obesity in humans might similarly lead to elevated OGT levels in TNBC tumors, which we noted from our OGT analysis of The Cancer Genome Atlas breast cancer subtype dataset (**Fig 1A-B**). If this pathway is true, it might be applied to explain higher levels of OGT expression in other forms of cancer,^54^ though we have not yet pursued this avenue of inquiry.

Inhibiting OGT might be one way to treat TNBC, as others have suggested.^27,28^ However, our OGT inhibition data for TARDBP production showed less of an effect at high glucose culture conditions compared to normal glucose conditions (**Fig 2C** vs. **Fig 5D**). Hyperglycemia leads to higher OGT substrate production (UDP-GlcNAc) in cells, and such elevated glucose to UDP-GlcNAc flux may outcompete the active site OGT inhibitor OSMI-4 in TNBC cells. If this finding is general, pre-clinical and clinical studies involving OGT may benefit from considerations of diet and physiology to account for elevated OGT levels and/or substrate availability in hyperglycemic settings. Also, effects of global OGT reduction might present untoward side effects because OGT is found in all mammalian cells.^25^ The mechanism presented here suggests downstream targets, including TET1,^36^ as potential therapeutic alternatives to OGT inhibition.

The cellular studies conducted in this report strongly suggest that hyperglycemia impacts CSC induction in TNBC via OGT, but there are several key limitations to this current study. Hyperglycemia is only one aspect of obesity/metabolic disease, so a more in-depth study of OGT-CSC pathway is needed using *in vivo* models that incorporate stable knockdown OGT/TNBC lines and/or OGT inhibitors with enhanced *in vivo* activity.^55,56^ Further studies are also needed to confirm that the OGT and O-GlcNAc effects we observed in obese vs. lean mice are also at play in patient tumors of various body mass indices. Finally, the impact of OGT inhibition was noticeably reduced in TNBC cells exposed to hyperglycemic conditions. The reduced impact of the OGT inhibition in hyperglycemic settings is a potential impact on obesity/OGT inhibitor effectiveness that will bear further examination.

The current report describes evidence to support that OGT expression level and activity drives a tumorigenic pathway in TNBC. An interesting facet of this pathway is “feed-forward” nature, whereby hyperglycemia leads to enhanced activity as well as elevated levels of OGT via the splicing factor TARDBP, located downstream of OGT in our model. Initial data in mice suggest that OGT is upregulated in DIO animals vs. lean littermates, potentially addressing one of the missing links between metabolic disease and TNBC. The majority of TNBC patients are obese,^49,50^ increasing the relevance of this study toward determine a potential mechanism for this obesity-TNBC correlation. Furthermore, obesity rates continue to rise globally.^57^ We envision that continued studies of OGT and O-GlcNAc-associated mechanisms in will help address a obesity-associated cancers as a critical unmet challenge in oncology.

## Authors’ Contributions

**Conception and design:** A. Bollig-Fischer, C. Fehl

**Development of methodology:** S. Ayodeji, B. Bao, A. Bollig-Fischer, C. Fehl

**Acquisition of data:** S. Ayodeji, B. Bao, E.A. Teslow, A. Bollig-Fischer

**Analysis and interpretation of data (e.g**., **statistical analysis, biostatistics, computational analysis):** S. Ayodeji, B. Bao, G. Dyson, A. Bollig-Fischer, C. Fehl

**Writing, review, and/or revision of the manuscript:** S. Ayodeji, A. Bollig-Fischer, C. Fehl

**Administrative, technical, or material support (i.e**., **reporting or organizing data, constructing databases):** S. Ayodeji, G. Dyson, A. Bollig-Fischer, C. Fehl

## Acknowledgments

This study was supported by NIH R35GM142637-01, awarded to CF. This study was also in part supported by funding from The Believe Foundation and Joann M. Deliz and James W. Deliz in memory of David Bergman (to ABF). The NIH:NCI Cancer Center Grant P30CA022453 to the Karmanos Cancer Institute supported contributions from the Biostatistics, AMTEC (Animal Model and Therapeutics Evaluation Core), and the MICR (Microscopy, Imaging, and Cytometry Resources) Cores. Mice used in this study were used from a previous study funded by these two former mechanisms. We thank the Suzanne Walker Lab (Harvard Medical School) for the kind gift of OGT inhibitor OSMI-4. pcDNA3-Flag-Tet1 CD was a gift from Yi Zhang (Addgene plasmid # 70129).

## Supplemental Data [duplicated from the separate “Supporting Information” .docx file]

**Supplemental Figure 1:**
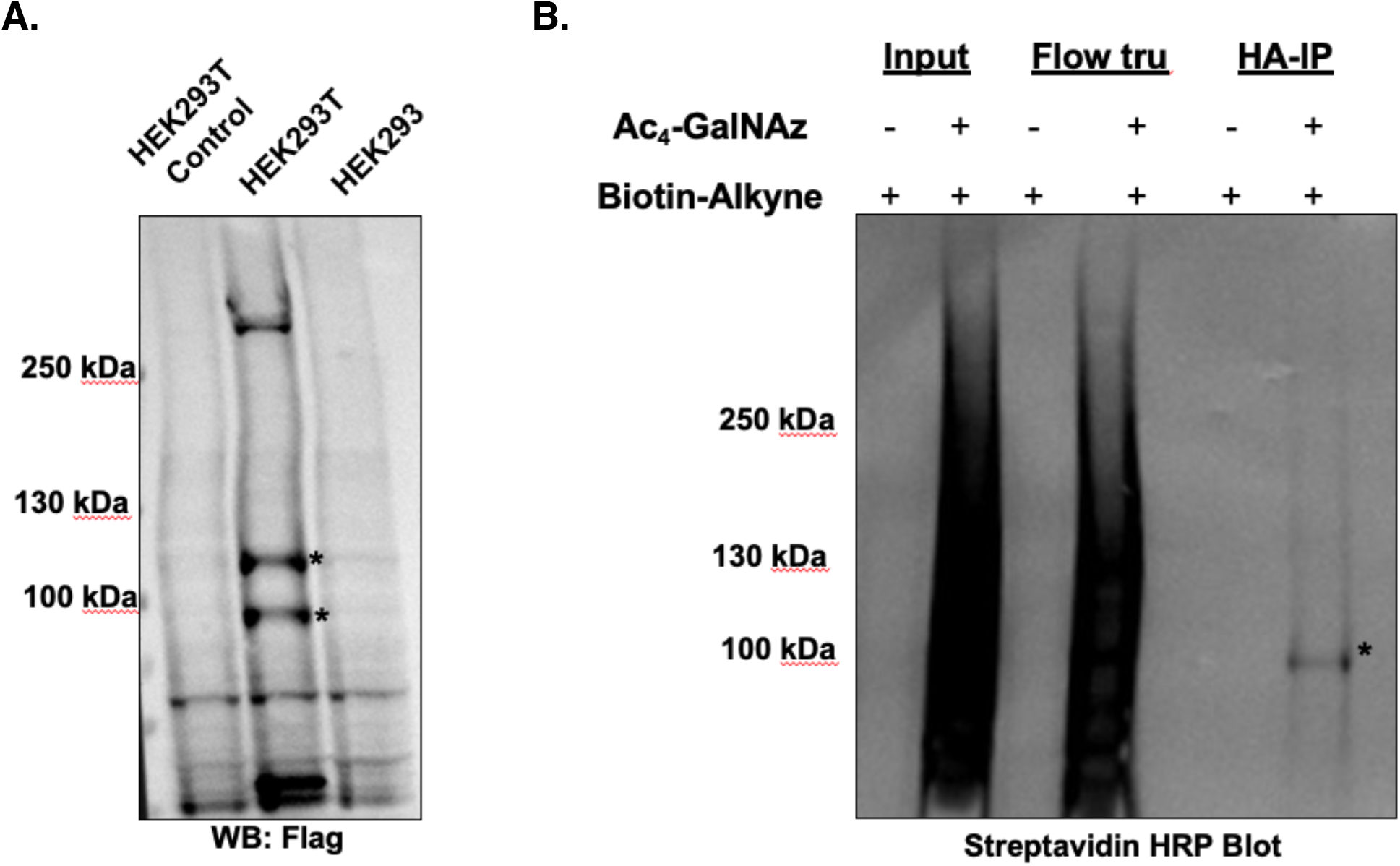
Alternative confirmation that TET1 is O-GlcNAc modified. **A**) Overexpression of Flag-HA-tagged TET1 catalytic domain in HEK293T and HEK293 cells. Stars indicate TET1-specific bands. **B**) HEK293T cells were fed the azide-labeled per-acetyl galactosamine (Ac4-GalNAz) as a metabolic reporter of O-GlcNAc. Click chemistry was performed with biotin alkyne. HA immunoprecipitation was used to enrich Flag-HA-TET1. Streptavidin blot was used to visualize biotinylated samples (O-GlcNAc proteins).

**Supplemental Figure 2:**
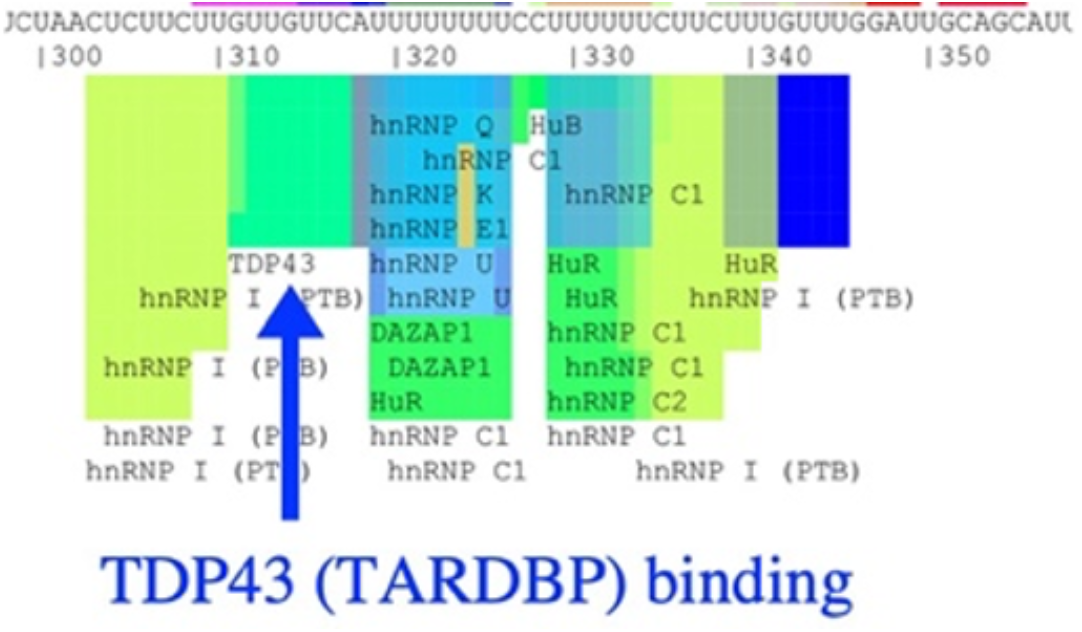
SpliceAid diagram of TARDBP binding site on OGT mRNA. Made using human OGT sequence and http://www.introni.it/splicing.html.

**Supplemental Figure 3:**
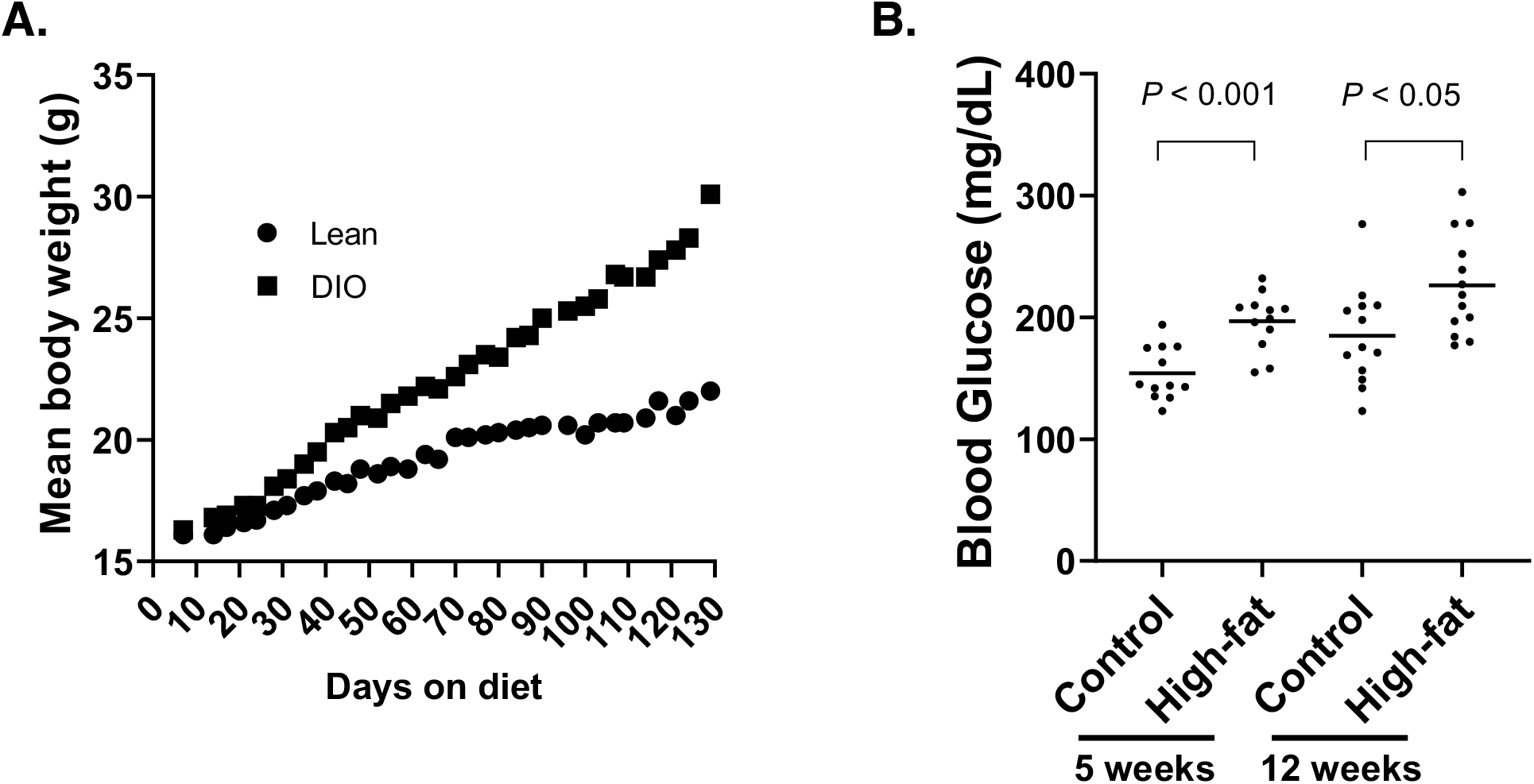
Increase in weight and blood glucose levels in female B6.Rag1-/-mice fed obesity-inducing high fat diet (60% fat) relative to control diet (10% fat). **A**) Weight gain over time. The difference was significant at 5 weeks, P < 0.001. **B**) Glucose in blood samples, compared control and high-fat diet at 5 and 12 weeks. 12 per group.

**Supplemental Figure 4:**
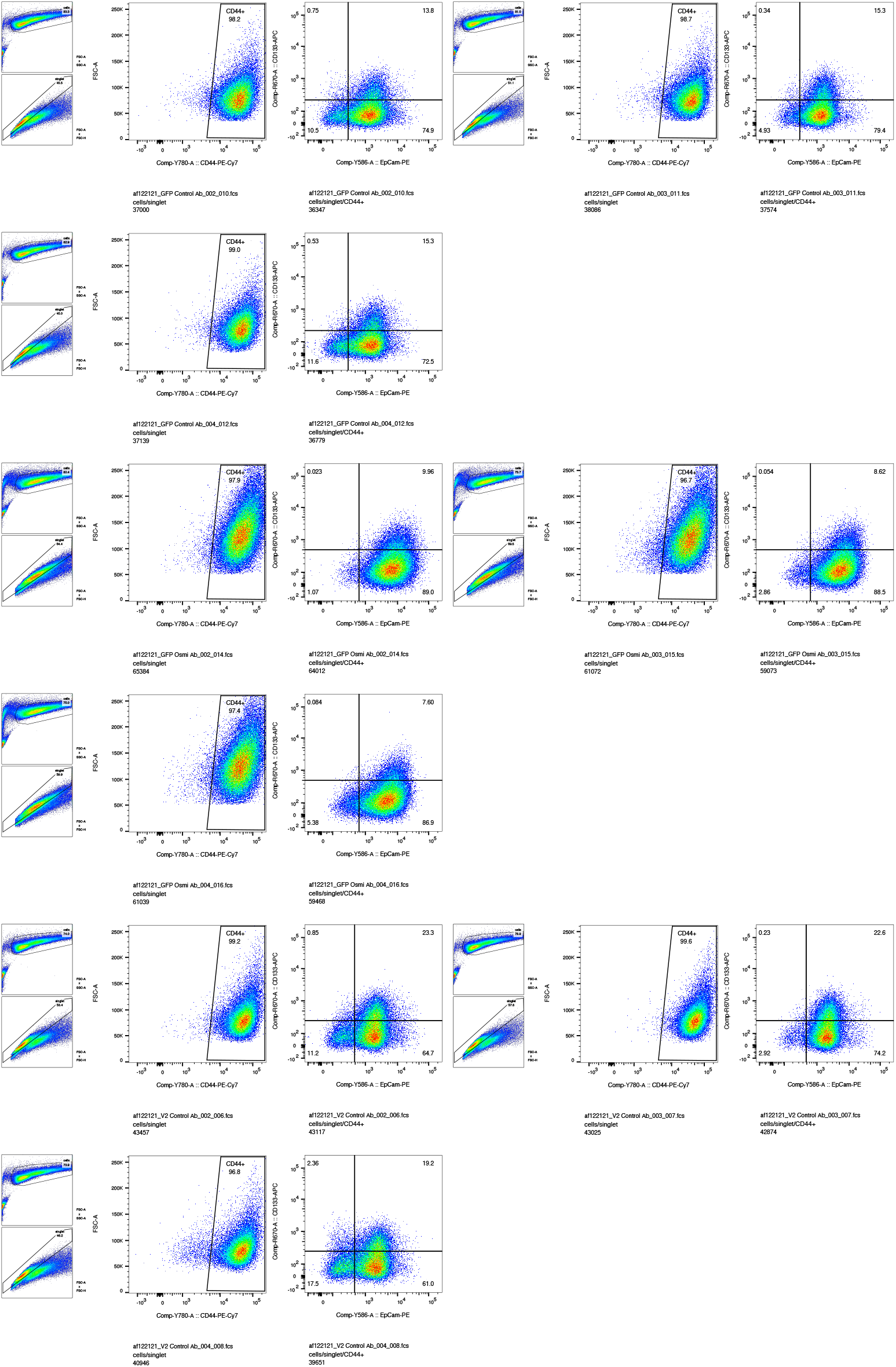
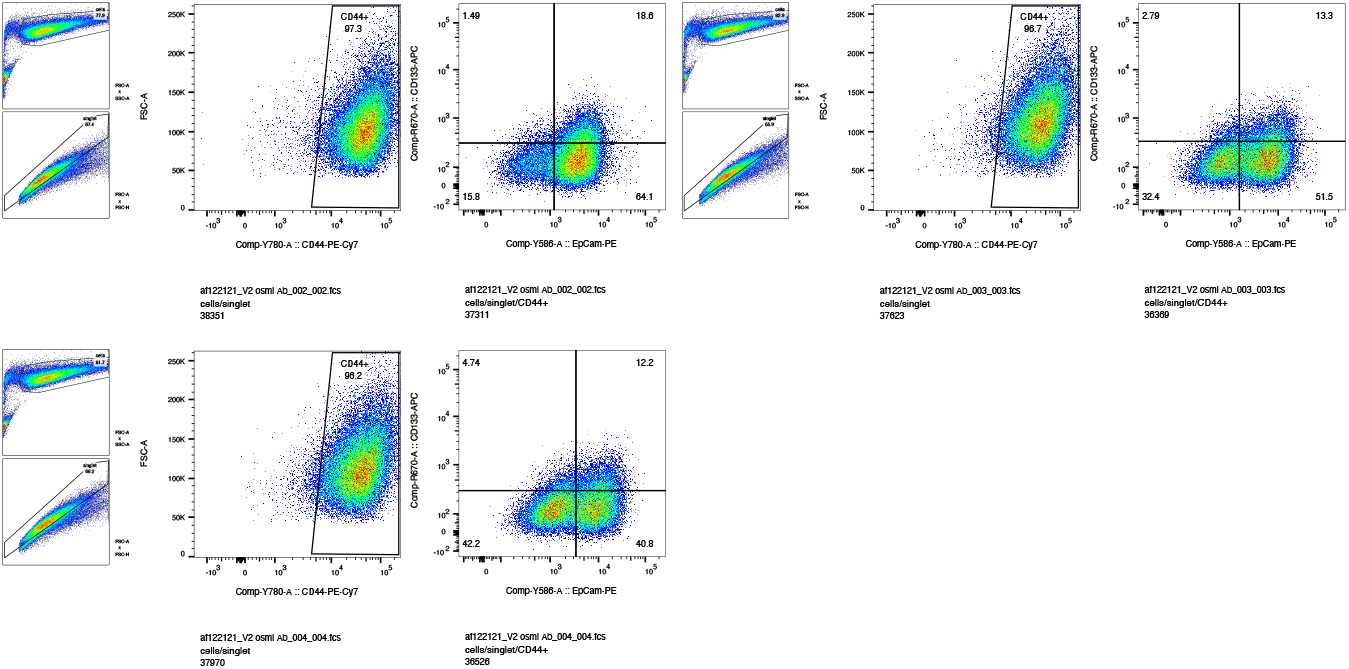
Analysis of cancer stem-like cell markers CD44, CD133, and EpCamAM in bulk TNBC cells. Cell lines: GFP-vector-expressing MDA-MB-468 cells or MBD2_V2-overexpressing MDA-MB-468 cell lines were treated with OGT inhibitor (OSMI-4, 10 uM, 24 h) and compared to untreated control. Cells were first gated by CD44, then analyzed for CD133 and EpCam staining. All three markers represent cancer stem-like cells. Results presented for three biological replicates per condition. FSC = front scatter plot and SSC = side-scatter plot, shown for reference to the left of all treatment conditions.

